# Evidence for compensatory evolution within pleiotropic regulatory elements

**DOI:** 10.1101/2024.01.10.575014

**Authors:** Zane Kliesmete, Peter Orchard, Victor Yan Kin Lee, Johanna Geuder, Simon M. Krauß, Mari Ohnuki, Jessica Jocher, Beate Vieth, Wolfgang Enard, Ines Hellmann

## Abstract

Pleiotropy, measured as expression breadth across tissues, is one of the best predictors for protein sequence and expression conservation. In this study, we investigated its effect on the evolution of cis-regulatory elements (CREs). To this end, we carefully reanalyzed the Epigenomics Roadmap data for nine fetal tissues, assigning a measure of pleiotropic degree to nearly half a million CREs. To assess the functional conservation of CREs, we generated ATAC-seq and RNA-seq data from humans and macaques. We found that more pleiotropic CREs exhibit greater conservation in accessibility, and the mRNA expression levels of the associated genes are more conserved. This trend of higher conservation for higher degrees of pleiotropy persists when analyzing the transcription factor binding repertoire. In contrast, simple DNA sequence conservation of orthologous sites between species tends to be even lower for pleiotropic CREs than for species-specific CREs. Combining various lines of evidence, we suggest that the lack of sequence conservation for functionally conserved pleiotropic elements is due to compensatory evolution within these large pleiotropic CREs. Furthermore, for less pleiotropic CREs, we find an indication of compensation across CREs. This suggests that pleiotropy is also a good predictor for the functional conservation of CREs, but this is not reflected in the sequence conservation for pleiotropic CREs.

## Introduction

One of the initial perplexing revelations of the human genome project was the seemingly limited number of genes, which did not align with the increase in complexity compared to organisms such as yeast, worms, and flies. It became evident that this complexity must stem from gene regulation, with the probability that most genes play roles in multiple contexts throughout development and in various tissues.

Considering the varying contexts of utilization in terms of location as well as timing, it follows that mutations within the same gene can exert influence on multiple traits. This phenomenon is widely recognized as pleiotropy. In a molecular context, pleiotropy is frequently measured as the number of tissues in which a gene is expressed, a metric called expression breadth (Hastings 1996; Duret and Mouchiroud 2000).

The advent of microarrays and subsequent RNA-seq technology allowed for an impartial, genome-wide evaluation of expression breadth. As data accumulated, it became evident that expression breadth is in fact a very good predictor of the conservation of protein sequences. In particular, the ratio of the non-synonymous over synonymous substitution rate (*d*_*a*_*/d*_*s*_) shows that pleiotropic genes tend to be more conserved than tissue-specific genes (Hastings 1996; Duret and Mouchiroud 2000; Zhang and WH Li 2004). Moreover, the amount of constraint added varies across tissues: Genes expressed in the brain tend to be more conserved than genes specific to other tissues, such as the liver (Kuma et al. 1995; HY Wang et al. 2007; Khaitovich et al. 2005). A similar pattern emerges in terms of expression level conservation; also brain-expressed as well as pleiotropic genes tend to have more similar expression levels across species than other genes (Khaitovich et al. 2005; Brawand et al. 2011; ZY Wang et al. 2020).

Naively, one would expect that a higher level of conservation of expression levels would be achieved via a higher level of conservation of the sequences of cis-regulatory elements (CREs). The resulting expectation would be that, if the same relationship between conservation and pleiotropy also applies to CREs and thus that CREs active in multiple tissues are also more conserved. However, most enhancers are tissue-specific (Gasperini et al. 2020) and show little conservation across species, although target gene expression appears conserved (Villar et al. 2014; Berthelot et al. 2018). Using a rather stringent definition of pleiotropy, a selection of a couple of hundred highly active pleiotropic enhancers was previously identified in humans and was found to have higher sequence conservation than tissue-specific enhancers across a large phylogeny (Andersson, Gebhard, et al. 2014; Singh and Yi 2021) and also over a much shorter evolutionary time scale focusing on genomic data from the human population (Huang et al. 2017).

Promoters are much more likely to be functionally conserved than enhancers (Berthelot et al. 2018). In addition, promoters are more pleiotropic than enhancers, which is probably due to the fact that core promoters are more restricted in their spatial genomic location than enhancers which can be located megabases away from the targeted transcription start sites (TSS). Promoters are further distinguished by their shape: Broad promoters are large, thought to harbor multiple TSS and tend to be more pleiotropic. In contrast, narrow promoters are small, probably have only one TSS and are more likely to be tissue-specific (Andersson and Sandelin 2020). Furthermore, evidence suggests that expression from broad promoters is less noisy and more robust towards mutations (Carninci et al. 2006; Schor et al. 2017; Sigalova et al. 2020; Floc’hlay et al. 2020) and in humans these broad promoters also show strong enrichment for CpG islands (Morgan and Marioni 2018). At least in flies, this results in the counter-intuitive observation that although broad promoters are more robust and thus also more likely to be functionally conserved across species, overall they exhibit lower sequence conservation between species than narrow promoters (Schor et al. 2017). In summary, the relationship between pleiotropy and sequence conservation for CREs appears to be much more complicated than that between pleiotropy and coding sequence conservation.

Here, we investigate the impact of pleiotropy on sequence and functional conservation in primates. To gauge pleiotropy, we thoroughly re-analyzed DNase hypersensitivity data from 9 primary fetal tissues (Bernstein et al. 2010), integrating across a minimal number of replicates to also identify tissue-specific CREs robustly. To assess functional conservation of the identified CREs, we obtained RNA-seq and ATAC-seq data from two human and two cynomolgus macaque neural progenitor cell lines. Furthermore, we obtained four different measures of sequence conservation: 1) a population genomic measure, 2) a conservation measure for the human lineage since the most recent common ancestor of humans and chimpanzees (Gronau et al. 2013), 3) a conservation score calculated for the primate phylogeny (Pollard et al. 2010) and 4) a scaled measure of transcription factor binding site (TFBS) conservation.

## Results

In order to investigate different aspects associated with varying degrees of regulatory pleiotropy, we identified putative CREs as DNase hypersensitive sites (DHS) in the Roadmap Epigenomics Data, which provide comparable experiments for a wide selection of tissues (Bernstein et al. 2010). To ensure reproducibility, we included only tissues for which at least seven biological replicates of DNase-seq data were available, leaving us with nine tissues: adrenal gland, brain, heart, kidney, large intestine, lung, muscle, stomach and thymus (Fig. 1A,B). We called DHS for each tissue separately using a peak caller that utilizes replicate information to gauge certainty (Ibrahim et al. 2015), resulting in a total of *>* 1.1 million DHS ranging from ∼ 80, 000 sites detected in the large intestine to ∼ 175, 000 sites detected in the stomach (Fig. 1C). In analogy to how expression breadth has been used as a proxy for pleiotropy of genes, we merge overlapping DHS from different tissues and define the Pleiotropic Degree (PD) as the number of tissues in which we found a DHS, resulting in ∼ 460, 000 union CREs stratified by PD. We distinguish promoters and enhancers based on genomic distance, while we designate CREs within 2kb of an active annotated TSS (Gencode v.32) as promoters and all other CREs within 1Mb as enhancers (Fishilevich et al. 2017; McLean et al. 2010).

**Figure 1.**
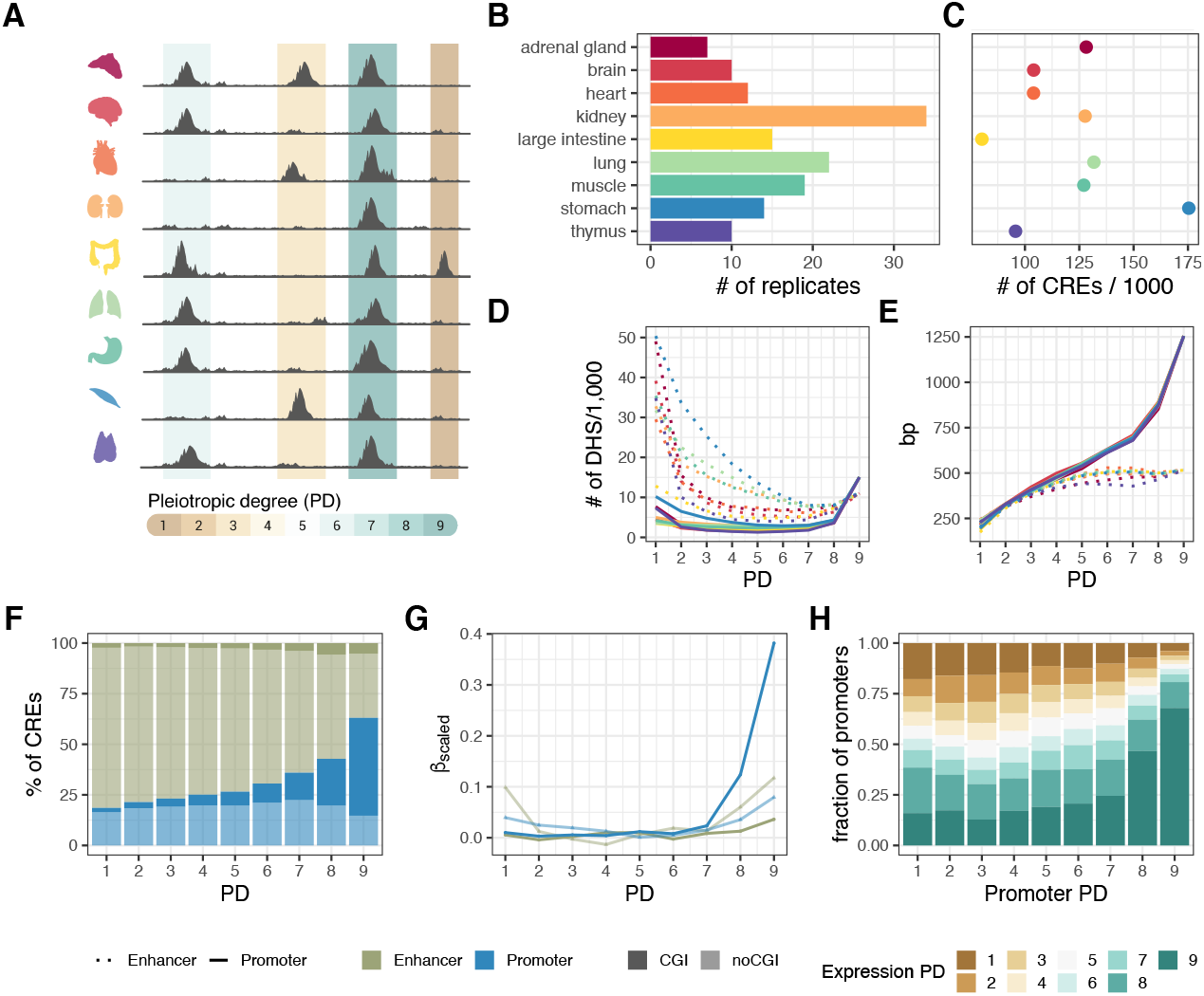
Study overview. (*A*) Open chromatin and expression data from the Roadmap Epigenomics Project (Bernstein et al. 2010) were used to infer the effect of pleiotropy on sequence and TFBS evolution, and associated gene expression in primates. Overlapping DHS peaks between tissues were merged to determine the degree of tissue-specificity per CRE. (*B* ) DHS-data from 9 human fetal tissues. The number of biological replicates per tissue varies between 7 and 34. (*C* ) The number of CREs per tissue varies 2.3-fold. There is no association between the number of replicates and the number of accessible regions per tissue, suggesting that with *>* 7 replicates per tissue, sufficient saturation is reached in peak detection. (*D* ) Most enhancers (dotted line) are tissue-specific, while promoters (solid line) are mostly pleiotropic. The colors represent the tissues as introduced in (*A,B* ). (*E* ) CRE length increases with the number of tissues, particularly at the promoters. This increase was also observed at the peak level prior to merging (Supplemental Figure S1A). (*F* ) The majority of PD9 CREs are CpG-island promoters (solid blue), while tissue-specific elements are rarely CpG-Islands and mainly enhancers (transparent green). (*G* ) Scaled coefficients of a linear mixed model to predict gene expression levels using distance scaled CRE counts of different types. (*H* ) Pleiotropic promoters are more commonly associated with pleiotropic gene expression patterns. The promoter PD indicates the highest PD of the associated promoters per gene. The y-axis shows the proportions of those x-categories (promoter PD) with associated gene expression pleiotropy ranging from 1 to 9.

Consistent with expectations, the majority of enhancers are tissue-specific (*PD*1) (Gasperini et al. 2020), while promoters are more likely to be pleiotropic (*PD*9) and CREs with an intermediate PD (1 *< PD <* 9) are rare among both promoters and enhancers (Fig. 1D). With a median size of 1.2kb, PD9 promoters are the largest CREs (Fig. 1E, Supplemental Fig. S1A) and also the overlap among the DHS inferred for each tissue is highest in PD9 promoters (Supplemental Fig. S1B), suggesting that their larger size is due to a higher content of information rather than being an artifact of concatenation. They probably correspond to the broad promoters observed in humans (Andersson, Gebhard, et al. 2014) and fruit flies (Schor et al. 2017). A large proportion of the pleiotropic promoters are CpG islands (76.7%) and the proportion of CpG island promoters generally decreases with increasing specificity (Fig. 1F). The same is true for enhancers, although enhancers are only very rarely CpG islands (3.2%). Next, we wanted to investigate whether the PD of a CRE has an impact on the expression of the associated genes. To this end, we integrated DNase-seq with gene expression estimates from matching samples that are also provided by the Epigenomics Roadmap Project (Supplemental Fig. S2). As expected, we find a strong enrichment for PD9 promoters to be associated with genes that are expressed in all 9 tissues, while we find an over-representation of tissue-specific promoters in tissue-specific genes (Fig. 1H). Moreover, we find that the pleiotropic degree of enhancers and promoters associated with a gene also has an impact on the gene’s expression level. The amount of variation in expression levels that can be explained by the number and distance of CpG island and non-CpG island CREs of varying PD is 24% (CI: 23.8-24.3%), while the number, distance and type of CRE without the pleiotropy information can only explain 19% (CI: 18.8-19.8%)(see Methods). Inspecting the scaled coefficients of the mixed effects model reveals that PD9 promoters have the largest activating effect on expression, followed by PD9 and PD1 enhancers. While for PD9 promoters, the signal is clearly due to CpG-island CREs, for enhancers the many non-CpG-islands CREs appear to have a larger activating effect in total (Fig. 1G; Supplemental Fig. S1C). We take this as evidence that our PD9 category as well as CREs that were found in only one tissue are likely to be functional.

### Characterization of transcription factor binding site repertoire across pleiotropic degrees

Under the premise that CREs regulate gene expression by binding transcription factors, we continued to characterize TFBS associated with CREs of varying pleiotropic degrees. To this end, we collected non-redundant position weight matrices (PWMs) of 643 binding motifs (Fornes et al. 2020) belonging to 561 TFs that we found to be expressed in at least one of the investigated tissues (Fig. 2A). Almost half of all expressed TFs (237 out of 561, 42%) were present in all tissues, i.e. pleiotropic, while 94 (17%) showed tissue-specific expression. Interestingly, we found that the brain has the highest proportion of tissue-specific TFs. Next, we evaluated the overall binding potential of a TF to a CRE using Cluster-Buster (Frith et al. 2003) (see Methods for details). Unsurprisingly, we found that TFBS diversity increases with pleiotropy for both enhancers and promoters. This is at least partially explained by the increase in CRE size, which is in turn likely linked to a broader functionality (Fig. 2B). Still, the question remains whether tissue-specific and pleiotropic CREs are regulated by the same TFs or whether preferences exist. For the majority of TFs we do not find a binding preference: 159 (24.7%) are over-represented in CREs specific for one of the tissues (Fig. 2C) and 84 (13.1%) motifs are enriched in the PD9 CREs. In line with our expectations, gene-set enrichment analysis shows that motifs enriched in brain-specific CREs are for TFs that are associated with neuron differentiation. Most prominently, this is driven by OLIG1 and OLIG2 that are essential for oligodendrocyte development (Zhou and Anderson 2002; Jakovcevski et al. 2009; Yu et al. 2013), as well as by NEUROD1, NEUROD2 and NEUROG1 that are important for neuron development (Olson et al. 2001; Sun et al. 2001; Messmer et al. 2012; Pataskar et al. 2016) (Fig. 2E). Other tissues also showed a specific enrichment: For example, TFBS that are overrepresented in heart-specific CREs include motifs of MEF2C, TBX20 and NKX2-5 (Fig. 2F), which are essential for cardiac muscle development (He et al. 2011; Schlesinger et al. 2011; Grunert et al. 2016).

**Figure 2.**
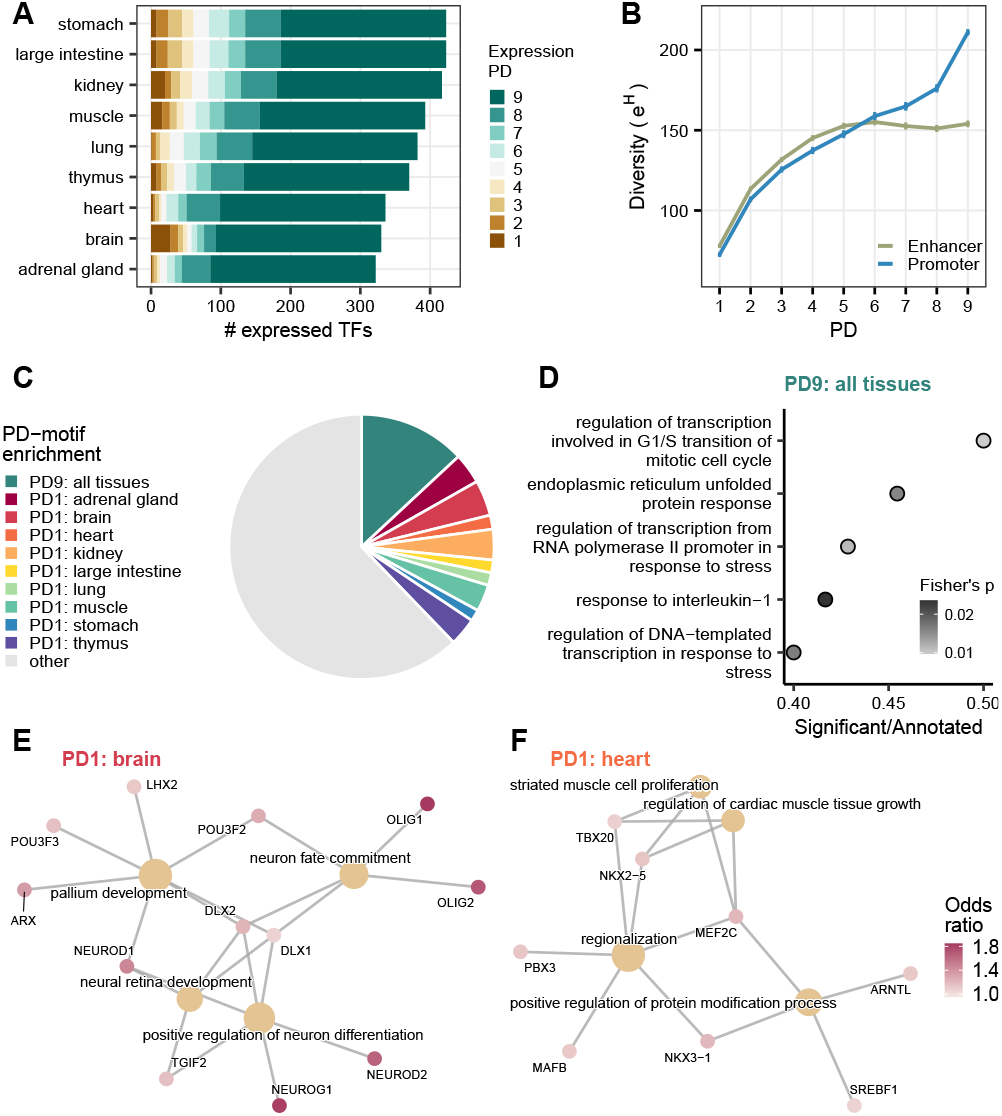
TFBS repertoire diversity and enrichment across tissue-specific and pleiotropic CREs. (*A*) An overview of TF expression across tissues. (*B* ) TFBS repertoire diversity increases with PD, particularly across promoters. Depicted are mean +/- SEM. (*C* ) Overview of the over-represented motifs in PD9 and PD1 CRE sequences. (*D* ) Top 5 categories of gene set enrichment analysis of PD9-enriched motifs using all motifs as background (Gene ontology, Biological Process, Fisher’s exact p-value*<* 0.05). (*E, F* ) Top 4 categories of gene set enrichment analysis of tissue-specific PD1 enriched motifs using all motifs as background (Gene ontology, Biological Process, Fisher’s exact p-value*<* 0.05). Fold change depicts the proportion of tissue-specific PD1 CREs with the motif over the global average proportion for that motif. (*E* ) Brain-specific PD1 over-represented motifs. (*F* ) Heart-specific PD1 over-represented motifs.

In contrast, TFs that show a binding preference for PD9 CREs appear to be associated with more basic cellular processes such as transcription regulation in connection with cell cycle and and stress response (Fig. 2D). These motifs are more GC-rich and tend to have a higher information content than PD1-enriched motifs or motifs without any preference (Supplemental Fig. S5A,B). In addition, these elements are enriched for TFs that were shown to co-localize with most other TFs (Odds Ratio = 10.51, Fisher’s exact p-value= 3*e* − 12) (Zhao et al. 2022). These so called “Stripe” TFs include SP, KLF and ZBTB family members, all of them recognize GC-rich sequences. Moreover, “Stripe” factors were experimentally shown to have a strong positive impact on prolonged CRE accessibility and the dynamics of most other TF proteins by stabilizing and prolonging their retention time at their binding site within the same CRE. Enrichment for binding sites for these universal and highly cooperative TFBS in PD9 CRE sequences is in line with the broad openness of these CREs and their high gene expression activating effects (Fig. 1G).

### The impact of pleiotropy on the evolutionary conservation of regulatory activity

To get a first glimpse of the interaction between the degree of pleiotropic and the evolutionary conservation of the CREs in our data, we generated RNA-seq and ATAC-seq data from iPSC-derived neural progenitor cell lines (NPCs) from humans and cynomolgus macaques (Supplemental Fig. S3A,B). We then intersected the detected genes and accessible peaks with the processed Epigenomics Roadmap data to assign a pleiotropic degree to the genes and peaks (Fig. 3A). As expected, the amount of CRE overlap with NPC ATAC-seq peaks increases with increasing PD and is generally higher for promoters than for enhancers (Fig. 3B). Moreover, the activity of PD9 CREs is also more conserved between humans and macaques. Of all the overlapping PD9 CREs, 88% were detected to be active in NPCs from both species, while this was only the case for 15% of the PD1 CREs. The observed dependence of PD on conservation levels is not only due to the increased activity that might generate a higher probability of PD9 elements being detected as peaks (Fig. 3 B). Instead, even without stratifying by whether a peak was called, we observe a decrease in differential activity with increasing PD, measured by absolute *log*_2_-fold changes (Fig. 3C).

**Figure 3.**
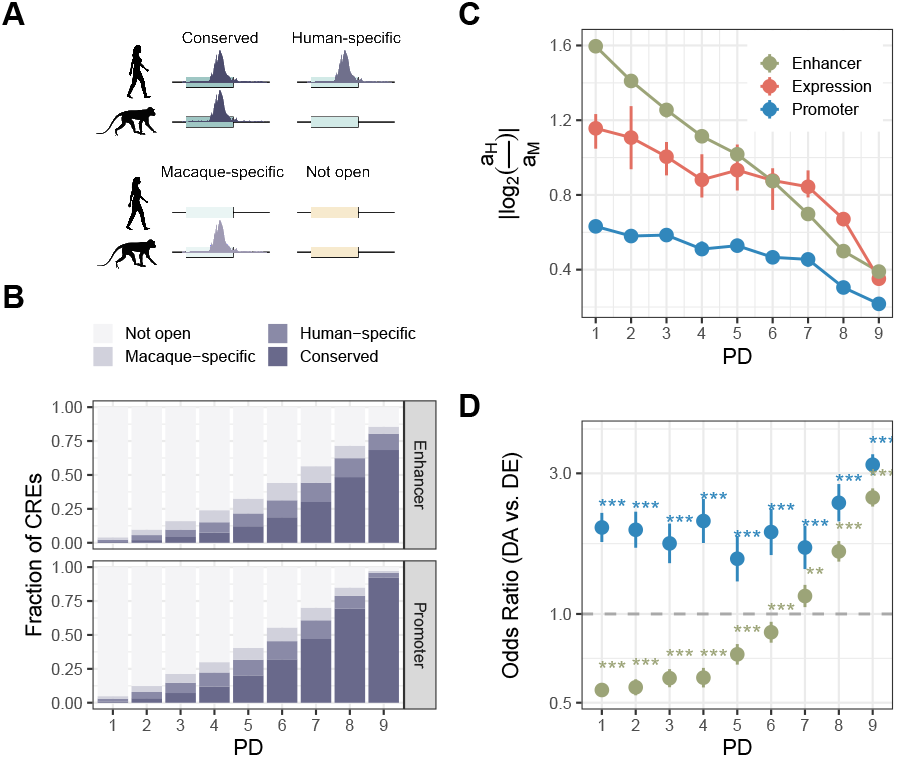
Pleiotropic degree and evolutionary conservation of expression and accessibility between humans and cynomolgus macaques. (*A,B* ) The fraction of enhancers and promoters of different pleiotropic degrees (PD) as defined using data from 9 tissues from the Epigenomics Roadmap project, which overlapped with ATAC-seq peaks called in neural progenitor cell lines (NPCs) from cynomolgus macaques and humans. The colors indicate whether a human DHS-derived CRE overlapped with a NPC ATAC-seq peak from humans, cynomolgus macaques, or both. (*C* ) Mean absolute log_2_-fold changes of gene expression and activities between humans and cynomolgus macaques. The error bars represent 95% bootstrap confidence intervals. PD9 genes (CREs) have more conserved expression (activity) than more tissue-specific genes. (*D* ) We tested for enrichment (odds ratio *>*1) or depletion (odds ratio *<*1) of differentially accessible CREs with significantly differentially expressed genes between humans and cynomolgus macaques. Error bars represent the 95% confidence intervals of the odd ratio, and the stars indicate the significance level with Benjamini-Hochberg correction ( . *<* 0.1, ^*^ *<* 0.05, ^**^ *<* 0.01, ^***^ *<* 0.001 ).

Next, we wanted to investigate which changes in CRE activity have an impact on the expression of the associated genes. To this end, we tested whether differentially accessible (DA) promoters and enhancers (BH-adjusted Wald test p-value *<*= 0.1) of a PD category are more likely to be associated with a differentially expressed (DE) gene (BH-adjusted Wald test p-value *<*= 0.1). Indeed, we find that DA promoters are more likely to be associated with a DE gene (BH adjusted Fisher’s exact test). There is a clear enrichment for all promoter PD categories, showing a 2-3 times enrichment (Fig. 3D). Moreover, when we further distinguish CpG island promoters, it turns out that the activity changes there have the greatest potential for downstream effects (Supplemental Fig. S3C,D).

This picture changes slightly for the association of enhancers: Although highly pleiotropic DA enhancers (PD8-9) are still more likely to be associated with a DE gene, for more tissue-specific DA enhancers, we observe a significant depletion in the associated DE genes (Figure 3D). In other words, genes with tissue-specific DA enhancers tend to have a more conserved expression. Generally, expression levels and robustness increase with increasing number of enhancers (Berthelot et al. 2018). Thus, if there are many enhancers, each has only a relatively small effect on expression and overall fitness, allowing these CREs to fluctuate between different possible genomic locations, resulting in different CREs for different species that can compensate for one another (Ludwig et al. 2000; Bradley et al. 2010; Doniger and Fay 2007; Arnold et al. 2014). In summary, the activity of pleiotropic CREs is evolutionarily more conserved between species than the activity of tissue-specific CREs. Moreover, if the activity of a pleiotropic CRE changes, such changes are also more likely to have downstream effects, i.e. to impact the expression of associated genes.

### Sequence conservation is lowest in pleiotropic CREs

So far, pleiotropy has the expected effect on gene regulation in that pleiotropic CREs tend to be more conserved. Here, we investigate how this functional conservation is reflected in the underlying DNA sequence. We focus on three measures of sequence conservation: 1) the number of weakly deleterious sites in humans (E.W. (Gronau et al. 2013), Fig. 4A,B), 2) the fraction of sites under (strong) negative selection (*ρ* (Gronau et al. 2013), Fig. 4C,D) and 3) the average phyloP and PhastCons scores across a primate phylogeny (Supplemental Fig. S4A,B) (Pollard et al. 2010). The main difference among the three measures is the evolutionary time across which sequence conservation is averaged. This ranges from recent selection within human populations (E.W.) via selection on the lineage since the most recent common ancestor of humans and chimpanzees (*ρ*), to the average across the primate phylogeny (phyloP, PhastCons). Since pleiotropic degree (PD) was assessed in human samples, the E.W measure provides the closest match to our measure of pleiotropy. For *ρ* and phyloP, we average the strength of selection over longer evolutionary times, and it is unclear whether the PD determined in humans has been constant. Additionally, it should be noted that variants emerging within a population may undergo recombination, whereas mutations occurring after speciation remain on separate haplotypes. In line with our expectations, we indeed find that the number of weakly deleterious sites increases with the pleiotropic degree for both promoters and enhancers (Fig. 4A). This observation aligns well with the conservation of CRE accessibility, which we assessed using the ATAC-seq data described above: Across all PD categories, we observe a higher prevalence of weakly deleterious sites in CREs that are open in both species (Fig. 4B). In contrast, when using *ρ* as a measure of conservation, we only find a higher sequence conservation for tissue-specific CREs (PD1-3) with conserved accessibility, while it appears that accessibility conservation is not reflected in the sequence conservation of pleiotropic CREs (Fig. 4D). Overall *ρ* suggests that PD9 CREs have the lowest fraction of negatively selected sites compared to other PD-categories (Fig. 4C). This surprising result remains when we use the average phyloP or average PhastCons score across a 10-species primate phylogeny as a measure of conservation, which confirms PD9 CREs as the PD category with the lowest conservation (Supplemental Fig. S4A,B). In summary, even though the number of weakly deleterious sites within a CRE increases with pleiotropy, this is not reflected in sequence conservation across species.

**Figure 4.**
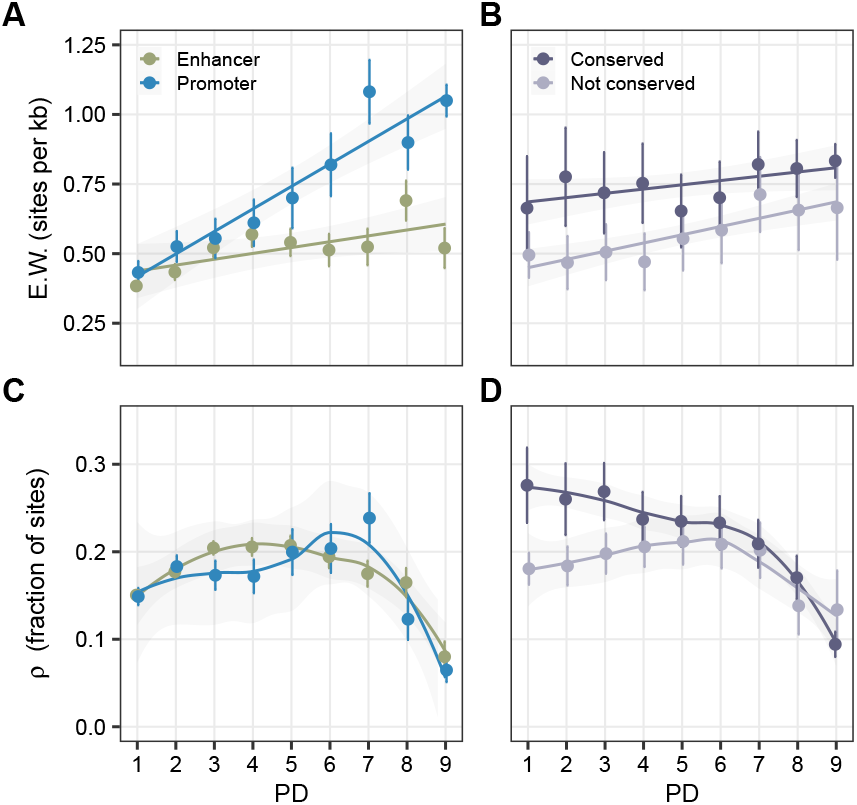
CRE sequence conservation patterns across varying degrees of pleiotropy. (*A,B* ) Weak negative selection inferred based on human polymorphisms increases with increasing pleiotropic degree (PD). (*A*) Separated by enhancers / promoters. (*B* ) Separated by human-macaque accessibility conservation in NPCs. (*C,D* ) (Strong) negative selection is the highest at the intermediately-specific CREs and lowest in the pleiotropic CRE sequences. (*C* ) Separated by enhancers / promoters. (*D* ) Separated by human-macaque accessibility conservation in NPCs. (*A,B,C,D* ) Depicted are mean estimates per PD category. Error bars indicate SEM.

### Tissue-specific effects

So far, we have not considered what happens if the different tissues would add different amounts of constraint. Indeed, when CREs are separated by the tissues in which they are utilized, the brain utilizes CREs that are clearly under more constraint than CREs of other tissues. Nevertheless, also for brain the number of weakly deleterious sites increases with PD, showing that although to smaller amounts, activity in other tissues still adds to the overall constraint (Fig. 5A). Again, this is not true when considering substitutions on the human lineage as used in the measure *ρ* (Fig. 5B). Here brain-specific CREs show most constraint on the human lineage, much more than pleiotropic PD9 CREs, which are by definition also utilized in the brain.

**Figure 5.**
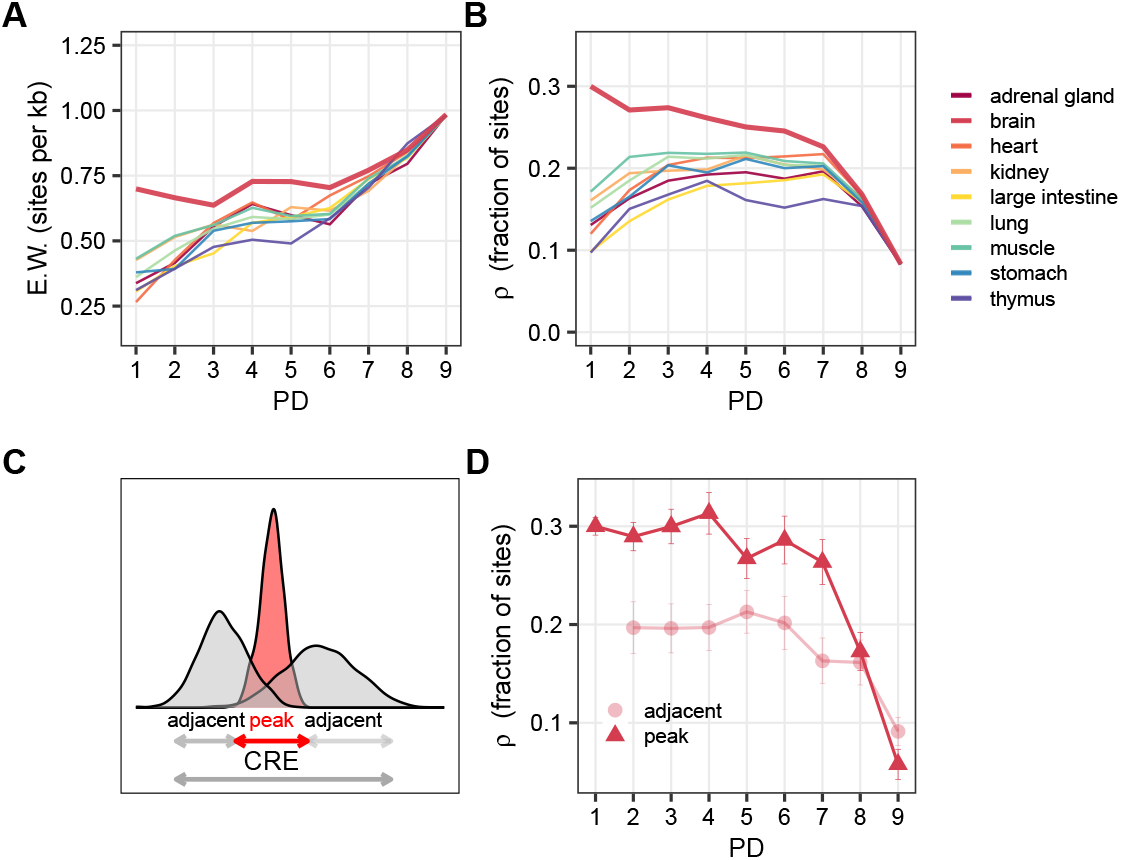
CRE sequence conservation patterns per tissue. (*A*) Weak negative selection inferred based on human polymorphisms separated by the tissue that utilizes the CREs. (*B* ) All negative selection separated by the tissue that utilizes the CREs. (*C* ) Brain CRE sequences, which showed the highest conservation across tissues, were separated into peak and adjacent sequences. (*D* ) The part of the sequence that is used by the brain shows much higher fraction of sites under negative selection than the respective adjacent sequences. (*A,B,D* ) Depicted are mean estimates per PD category. Error bars indicate SEM.

To exclude the possibility that the brain effect on the PD9 elements is diluted by the merging of DHS across tissues, we contrast the *ρ* of the brain peak sequence with adjacent sequences that are part of the same merged CRE but are open in other tissues (Fig. 5C). We find that for PD9 CREs, brain peak sequences show lower sequence conservation on the human lineage than the adjacent sequence utilized only by other tissues, while for less pleiotropic CREs the part that is used in the brain is under much more constraint (Fig. 5D). In summary, even though we find tissue-specific effects, in particular a higher constraint for brain CREs, this cannot explain the overall pattern of the relatively low sequence conservation of pleiotropic CREs. It remains that for pleiotropic CREs there is no simple relationship between sequence and functional conservation between species.

### Pleiotropic CRE TF repertoire is conserved, not the binding sites

In order to explain the apparent mismatch between functional and sequence conservation in PD9 CREs across primates, we continued to analyze levels that are intermediate between sequence conservation (less functional) and accessibility conservation (more functional), which are CpG content, TFBS repertoire and position conservation between human CREs and their orthologous sequences in cynomolgus macaques. To begin with, we find that conservation of CpG content increases with PD and is highest for pleiotropic promoters (Supplemental Fig. S4E). This coincides with the increase in CpG island CREs with PD (Fig. 1F) and suggests that the CpG island properties are conserved across species, landing closer to the functional side. Next, we calculated the binding potential for all expressed TFs and calculated the average pairwise Canberra distance between species 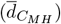. We then approximate TFBS repertoire conservation as 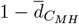 . To ensure that repertoire conservation is not dominated by differences in diversity between PDs, we shuffled the CRE identifiers of the macaque profiles within the respective PD class and calculated the average random TFBS profile similarity between species (Supplemental Fig. S5C,D). Furthermore, when we contrast CREs with conserved and non-conserved openness between humans and macaques, we find that for all PD categories, functionally conserved CREs also show a higher repertoire conservation (Fig. 6B).

**Figure 6.**
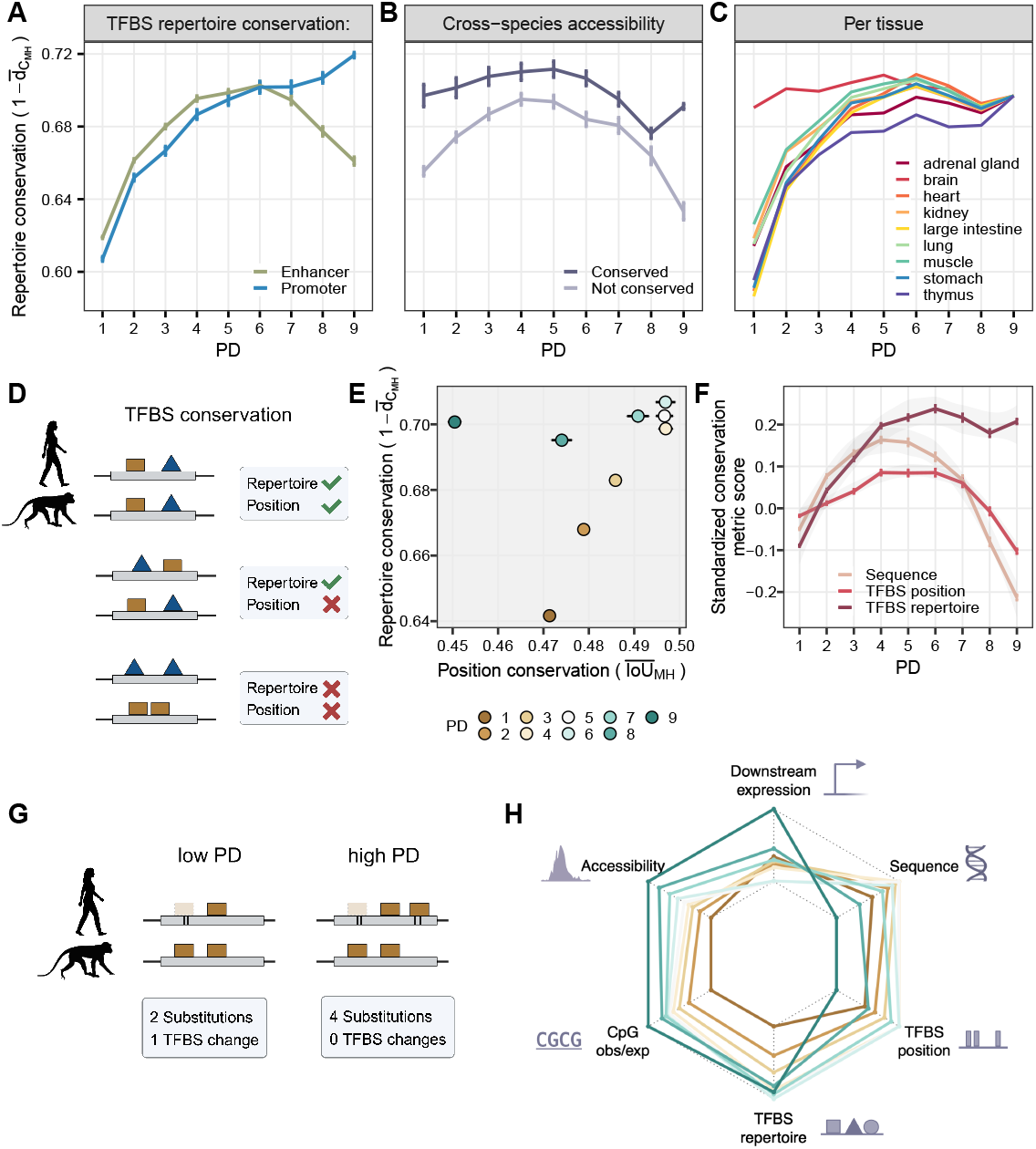
TFBS repertoire and position conservation between orthologous human and macaque CREs. (*A,B,C* ) TFBS repertoire conservation across PDs. Depicted are mean +/- SEM. (*A*) TFBS repertoire conservation increases with higher PD among promoters, however, it decreases slightly at high PD-enhancers. (*B* ) CREs that overlap NPC peaks with conserved openness show higher TFBS repertoire conservation than species-specific NPC peaks. (*C* ) TFBS repertoire conservation differs across tissues, where brain shows the highest conservation at lower PDs. (*D* ) Simplified schematic of the measures of repertoire and position conservation. (*E* ) TFBS position conservation versus repertoire conservation across PD categories. Depicted are mean values +/- SEM. (*F* ) Standardized scores (z-scores) of sequence (primate phyloP), TFBS repertoire and binding site conservation between human and cynomolgus macaque. (*G* ) A schematic depicting how lower sequence conservation might lead to higher TFBS repertoire conservation through compensatory mechanisms. (*H* ) A summary of the scaled average conservation metric scores across PDs. Sequence: primate phyloP scores, TFBS position: 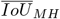 scores, TFBS repertoire: 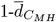, CpG observed/expected: 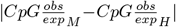, accessibility: —LFC— of NPC-DA results, downstream expression: —LFC— of NPC-DE.

With respect to the PD categories, we found that repertoire conservation generally increases with pleiotropy in all tissues (Fig. 6A,C). However, while there is a simple relationship for promoters for which repertoire conservation is highest for PD9 and lowest for PD1 CREs, this is not the case for enhancers among which CREs with intermediate PDs show the highest conservation. This said, also for enhancers repertoire conservation in PD9 CREs (0.66) is considerably higher than PD1 TFBS repertoire conservation (0.62), which is in contrast to what we observed for sequence conservation, again showing overall a higher similarity to the functional pattern of conservation (Fig. 3D, 6F). To answer in more detail how for PD9 CREs a relatively high repertoire conservation is achieved in spite of a low sequence conservation, we analyzed the positional conservation of TFBS as a third intermediate metric. We calculated the per-motif position conservation as the fraction of conserved binding sites between both species (intersection) over the total binding sites per motif across species (union) (Jaccard similarity index 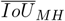) (Fig. 6D). Surprisingly, we find that the average repertoire conservation appears to be unrelated to the positional conservation in high PD categories (Fig. 6E). The positional conservation seems to be more related to sequence conservation, thus landing on the less functional side (Fig. 6F). In summary, while CRE sequence and TFBS positions are least conserved in PD9 elements, CpG content and TFBS-repertoire are in agreement with the more functional metrics accessibility and expression conservation in that they show the highest conservation in PD9 elements. These puzzling patterns would be consistent with a mechanism of compensatory evolution. In a simplified scenario, if a certain TF binding site is lost in a more tissue-specific CRE and no new binding site is fixed to compensate for this, this would lead to fewer substitutions than in the case where the loss of a binding site is compensated by the fixation of a new binding site (Fig. 6G). Such compensation in the latter case would lead to a low sequence and positional, but high repertoire conservation. Many genome-wide studies have confirmed that TFBS have a high turnover rate (Dermitzakis and Clark 2002; Paris et al. 2013; Domené et al. 2013), which is buffered by compensation. Here, we describe the evolutionary patterns where this compensation likely happens within the same CRE.

### PD9 promoter of Ataxin-3 gene as an example

To illustrate within CRE compensatory evolution of TFBS within a PD9 promoter, we took a closer look at the promoter of the ubiquitously-expressed protein-coding gene ATXN3 (Ataxin-3). Ataxin-3 is an important factor for the regulation of the degradation of damaged proteins (Schmitt et al. 2007; Gao et al. 2015; Feng et al. 2018). This gene plays an important role for the brain, as its malfunction can lead to neurodegenerative diseases such as spinocerebellar ataxia (Evers et al. 2014). The ATXN3-promoter shows low sequence conservation (34%) and low TFBS binding site conservation (49%), but high TFBS repertoire (77%), accessibility and expression conservation (Fig. 7A-E).

**Figure 7.**
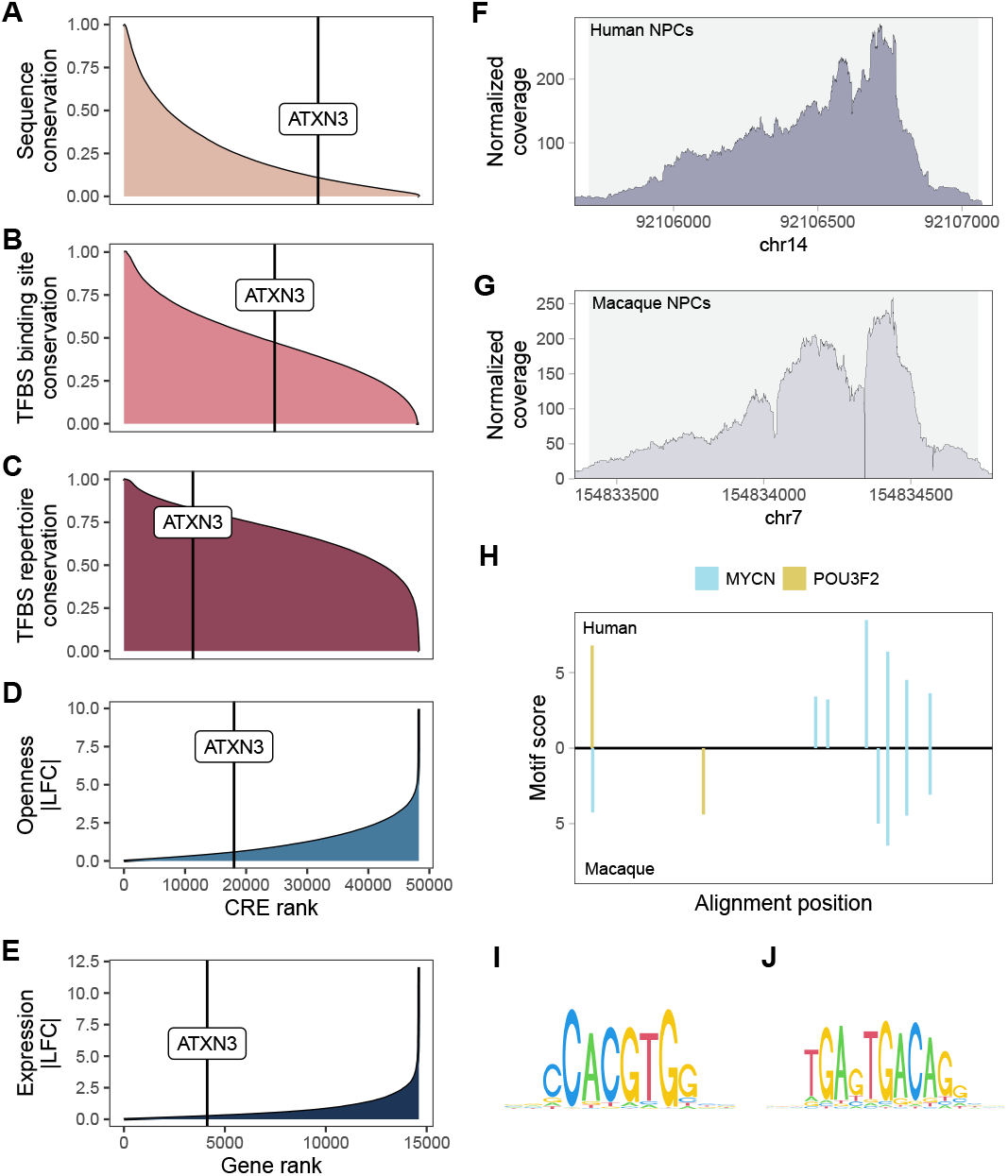
Ranks of ATXN3 PD9 promoter compared to other CREs in terms of (*A*) sequence conservation (mean PhastCons), (*B* ) TFBS binding site conservation, (*C* ) TFBS repertoire conservation, (*D* ) CRE openness conservation between human and cynomolgus macaque in NPCs and (*E* ) ATXN3 gene expression conservation between human and cynomolgus macaque in NPCs. (*F, G* ) ATXN3 PD9 promoter is accessible in both species. (*H* ) ATXN3 promoter shows diverged TFBS positions between species among validated TFs involved in neurogenesis. (*I, J* ) PWM logos of the investigated TF motifs with ChIP-seq data available: MYCN (*I* ), POU3F2 (*J* ).

To investigate a few likely relevant TFs closer, we overlapped our TFBS data with published ChIP-seq data from human neural cells available in the GTRD database (Yevshin et al. 2018) and visualized the binding sites of the 2 TFs (MYCN, POU3F2) annotated to be involved in neurogenesis (Gene Ontology biological process term GO:0022008) (Fig. 7H). Both of their motifs are moderately complex as shown by their information content (MYCN: IC=11.8, POU3F2: IC=13.7, Fig. 7I,J). Both promoter orthologues show strong binding positions for both TFs. Humans have 6 and macaques 5 MYCN binding sites and both have one POU3F2 binding site, suggesting a rather high repertoire conservation, which is also reflected in similar ATAC-seq peak-shapes (Fig. 7F,G). However, only 3 of the 10 binding sites are positionally conserved between the species. This serves as an example of how the large disagreement between sequence, TF binding site conservation and TFBS repertoire might co-occur.

## Discussion

Pleiotropy has been shown to be the best predictor of both protein coding sequence conservation (Hastings 1996; Duret and Mouchiroud 2000; Zhang and WH Li 2004) and gene expression levels (Khaitovich et al. 2005; Brawand et al. 2011; ZY Wang et al. 2020). Here, we investigate the effect of pleiotropy on the evolution of cis-regulatory elements (CRE) and find that measures close to CRE function, such as accessibility and TFBS repertoire conservation, indeed show the expected higher conservation for more pleiotropic CREs. Similarly, a measure of conservation based on human diversity data also shows a trend for higher conservation in more pleiotropic CREs. However, surprisingly, we found that this higher conservation of pleiotropic CREs is not reflected in the sequence and positional conservation of TFBS between macaques and humans. These observations imply that a simple model of purifying selection alone is insufficient to explain the effect of pleiotropy on CRE evolution and suggest a role for compensatory evolution.

Zooming into tissue effects, in line with previous investigations on brain evolution (Kuma et al. 1995; HY Wang et al. 2007; Brawand et al. 2011), we find that the activity in the brain exerts more constraint on a CRE than the activity in other tissues. There are many reasons why the brain is special and requires particularly tight regulation, including its high complexity consisting of precise neural networks (Geschwind and Rakic 2013). Hence, it comes as no surprise that brain-specific CREs show by far the highest sequence conservation irrespective of the measure. However, following the logic that brain expression induces a lot of constraint, this should also impact the pleiotropic, i.e. PD9 elements. Looking at the between-species sequence conservation measure, the sequences of PD9 CREs that are open in the brain are even less conserved than the adjacent sequences (Fig. 6D). This confirms the notion that the structure and evolution of PD9 CREs is inherently different, in that it allows for functional conservation without much sequence conservation.

Indeed, we find several basic structural properties of PD9 elements that distinguish them from less pleiotropic CREs. They tend to be larger, have more CpGs and a higher GC content. Moreover, PD9 elements show an over-representation of GC-rich motifs that are associated with TFs that tend to be involved in more basic cellular processes. Among those, we also find enrichment for binding sites of a recently described group of highly cooperative TFs (Universal Stripe factors) that prolong CRE openness (Zhao et al. 2022). In concordance with the idea that these Stripe factors facilitate the binding of most other TFs, we observe that PD9 CREs are more diverse in their TFBS. It should also be noted that the majority of PD9 CREs are promoters and PD9 promoters share many properties with broad promoters that were defined via the shape of CAGE peaks (Andersson, Gebhard, et al. 2014). Even though this classification is based on a completely different concept, also broad promoters were shown to be more pleiotropic, active and CpG-rich. Indeed, as observed for PD9 CREs, broad promoters also tend to show an increased substitution rate. Moreover, broad promoters have been shown to be more robust than narrow promoters, in that they show less expression noise across haplotypes in Drosophila (Floc’hlay et al. 2020; Schor et al. 2017). Similarly, in humans CpG island promoters have also been found to induce more stable expression (Morgan and Marioni 2018). Mechanistically, this picture fits nicely with the notion that Stripe factors bind to GC-rich regions, thus facilitating combinatorial binding, which has been shown to lead to evolutionarily more stable TF binding across mouse species (Stefflova et al. 2013). In the same vein, Hagai et al. 2018 found that the regulatory response of genes associated with CpG islands to an immune stimulus is more conserved than that of genes associated with a TATA-box. In summary, there is ample evidence that large CpG island promoters are functionally robust while having high substitution rates.

Nearly as pronounced as for promoters, we also find high substitution rates in PD9 enhancers, which also share most of the other features with PD9 promoters, suggesting that similar evolutionary mechanisms apply to both promoters and enhancers. We suggest that the main differences in the evolutionary patterns observed for promoters and enhancers are closely linked to the degree of pleiotropy. Most enhancers show strong tissue preferences, placing them in our PD1 category. Consistent with multiple other studies investigating CRE conservation in mammalian genomes (Danko et al. 2018; Berthelot et al. 2018), we find that only a relatively small fraction of enhancers is conserved between species in terms of accessibility and that these fractions show a strong association with the pleiotropic degree irrespective of their classification as enhancers or promoters. In fact, the CRE conservation across the genome is so puzzlingly low (Doniger and Fay 2007; Crocker et al. 2016; Horton et al. 2023), implying such high TFBS turn-over rates beyond what simple models of evolution can explain (Tuğrul et al. 2015).

In addition, the observed high TFBS turnover rates appear to be inconsistent with the relatively low rates of change in gene expression levels. This discrepancy has prompted the proposal of compensatory evolution as a prevalent mechanism for CREs. The phenomenon of CREs at non-orthologous genomic positions in different species exhibiting the same function and being able to compensate for one another has been documented for several cases (Ludwig et al. 2000; Arnold et al. 2014; Domené et al. 2013). Also in our data, we find hints that the between-CRE compensation impacts the evolution of cis-regulatory networks between humans and macaques. The positions of more tissue-specific enhancers appear to be less conserved for genes with conserved expression (Fig. 5D). This phenomenon is related to the observation that a lot of function is encoded redundantly also within a gene’s regulatory landscape by the so called shadow enhancers (Hong et al. 2008; Osterwalder et al. 2018; Wunderlich et al. 2016). Osterwalder et al. 2018 showed that the deletion of one strong enhancer did not have an effect on the phenotype as long as the shadow enhancer was still active. This clearly demonstrates the presence of epistasis, which suggests that multiple equally fit haplotypes exist and a different ones can get fixed in each species, which is then perceived as compensatory evolution across CREs.

Several other properties of CREs suggest that there is a lot of epistasis also within one CRE. The billboard model (Kulkarni and Arnosti 2003; Arnosti and Kulkarni 2005) and the TF-collective model (Junion et al. 2012) of enhancer activity suggest that two CRE haplotypes with shifted but similar TFBSs should be functionally equivalent. It follows that the mutations that create these two haplotypes will also have a non-additive effect on fitness. Moreover, some studies showed that binding to a high-affinity site is facilitated by many neighboring low-affinity binding sites, thus providing the raw material for high TFBS turnover rates (Tuğrul et al. 2015).

Thus, we suggest that within-element compensation of TFBS is a common mode of evolution for pleiotropic CREs. This would explain the apparent disparity between the cross-species sequence conservation and the within-species constraint measure E.W. (Fig. 4). Moreover, it would also explain the disparity between the low sequence and the high functional conservation between species as observed in our ATAC-seq and RNA-seq data: If different, functionally equivalent haplotypes got fixed in different species, this would lead to a high sequence divergence while the open chromatin state and downstream gene expression remained conserved (Fig. 6H). Furthermore, we show that even though PD9 TFBS may not have a high positional conservation, the overall binding potential for various TFs across a pleiotropic CRE tends to be conserved.

In summary, we think that compensatory evolution is a prevalent mode for evolution of regulatory elements and goes along with the number of contexts in which the element is utilized. The structure of cis-regulatory networks lends itself to high levels of negative epistasis across more distal CREs, while for the complex, large pleiotropic CREs epistatic interactions are more likely to occur within the same element. The within-element compensation is possibly facilitated by higher spatial restrictions on TFBS locations: Promoters are likely more restricted spatially than enhancers. However, we observe similar patterns for pleiotropic enhancers as well, albeit less pronounced. We speculate that they are also spatially more restricted than less pleiotropic enhancers due to their higher sequence complexity, which is probably due to highly cooperative binding at the pleiotropic sites. Such complex element structures are less likely to spontaneously occur at distal sites than it is observed for tissue-specific elements.

## Methods

### Human DNase-seq and RNA-seq data

DNase-seq and RNA-seq data from human fetal tissues of week 10-20 generated within the Roadmap Epigenomics project (Bernstein et al. 2010) were downloaded from the NCBI’s Sequence Read Archive (Dec. 15, 2014, summary table on github). We included only tissues for which at least 7 biological DNase-seq replicates from primary tissue samples were available. This left us with 9 different tissues: adrenal glands, brain, heart, kidney, large intestine, lung, muscle, stomach, and thymus.

### Cis-regulatory element (CRE) region determination and tissue-specificity scoring

DNase-seq reads were mapped to human genome version hg19 using NextGenMap (Sedlazeck et al. 2013, version 0.0.1). Aside from a few exceptions (dualstrand = 1; min identity = 0.9; min residues = 0.5), the default parameters were used. PCR duplicates were removed using samtools rmdup (H Li and Durbin 2009, version 1.1). We used JAMM (Ibrahim et al. 2015, version 1.0.7) to call peaks per tissue considering the biological replicates for the DNase-seq data using the recommended settings. To compare peaks across tissues, we merged overlapping peaks using the resulting union peaks as putative CRE, which are the basis of most further analyses. We removed peaks mapping to Y or MT chromosomes. Furthermore, we removed 26 CREs whose width exceeded 5000 bp (*<* 0.0001%), resulting in a set of 465,281 CREs. We then used the number of overlapping peaks, i.e the number of tissues in which a CRE is accessible as a proxy for pleiotropy. This score ranges between 1 (tissue-specific) to 9 (ubiquitously open).

### CRE annotation and association with genes

We used transcript annotation for hg19 from Gencode v.32 (Harrow et al. 2012) where we considered each transcript 5’ end as a transcription start site (TSS). For each tissue we only considered TSSs of the expressed genes in the complementary RNA-seq data. CREs within 2 kb of a TSS are designated as promoters and associated with all TSSs within that distance. All other CREs within 1 Mb of a TSS are deemed to be enhancers and are associated with the 2 closest TSSs (one in each direction), unless the distance to one TSS is at least 10x smaller than to the other TSS - in that case only the closest TSS is assigned. In total, we could assign 443,322 out of 465,281 CREs (95.3%).

### CRE effect on gene expression across tissues

For each of the included tissues, RNA-seq RPKM expression matrix was filtered to include only genes that are detected with *>*1 count in 50% of the samples in that tissue. Number of included genes varies from 12,283 (brain) to 19,382 (lung). Log mean expression was modeled as a linear mixed model with tissues as a random effect and the distance to TSS weighted (*d*) numbers of CpG Island and non-CpG Island promoters and enhancers that was fit using the lme4 function from the lmer package (version 1.1-30) in REML mode:

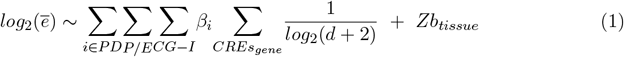

For the comparability, we report the standardized coefficients *β*_*scaled*_ = *βs*_*x*_*/s*_*y*_ and the marginal coefficient of variation as calculated for generalized linear mixed models was done with the R-package part2 (Nakagawa et al. 2017, version 0.9.1.9000). In order to assess the effect of PD independently of the distance and number of CREs, we shuffle the PD across all CREs while keeping all other parameters constant and calculate and compare those estimates.

### Human and cynomolgus macaque iPSC differentiation into NPCs

Previously generated urinary stem cell derived iPS-cells of 3 human individuals (*Homo sapiens*) and fibroblast derived cynomolgus macaque iPSCs (*Macaca fascicularis*) of 2 individuals (Geuder et al. 2021) were differentiated to neural progenitor cells via dual-SMAD inhibition as three-dimensional aggregation culture (Chambers et al. 2009; Ohnuki et al. 2014). Briefly, iPSCs were dissociated and 9x10^3^ iPSCs were seeded in a low attachment U-bottom 96-well-plate in 8GMK medium consisting of GMEM (Thermo Fisher), 8% KSR (Thermo Fisher), 5.5 ml 100× NEAA (Thermo Fisher), 100 mM Sodium Pyruvate (Thermo Fisher), 50 mM 2-Mercaptoethanol (Thermo Fisher) supplemented with 500 nM A-83–01 (Sigma Aldrich), 100 nM LDN 193189 (Sigma Aldrich) and 30 μM Y27632 (biozol). Culture medium of the spheres was changed every second day until they were harvested or plated for further culture. In order to obtain stable NPC lines, spheres were dissociated on day 7 of the differentiation process using Accumax (Sigma Aldrich) and plated onto Geltrex (Thermo Fisher) coated dishes. NPCs were subsequently cultured in NPC proliferation medium (DMEM F12 (Fisher Scientific) supplemented with 2 mM GlutaMAX-I (Fisher Scientific), 20 ng/mL bFGF (Peprotech), 20 ng/mL hEGF (Miltenyi Biotec), 2% B-27 supplement (50×) minus vitamin A (Gibco), 1% N2 supplement 100× (Gibco), 200 μM L-ascorbic acid 2-phosphate (Sigma), and 100U/ml 100μg/ml penicillin-streptomycin). All cell lines have been authenticated using RNA sequencing (RNA-seq) (Geuder et al. 2021), and the current study.

### RNA-seq data generation and processing

Samples for RNA-seq were taken from 3 clones of 3 human individuals and 4 clones of 2 cynomolgus macaque individuals at the iPSC stage (time point 0) and after 1, 5, 7 and 9 days during the neural maturation process. Spheres were dissociated at each time point using Accumax (Sigma Aldrich) and live cells were sorted using the BD FACS Aria II.

cDNA libraries for samples from the different species and differentiation time points were generated using the prime-seq protocol (Janjic et al. 2022) and we obtained 100bp cDNA reads from a Illumina HiSeq 1500 and another read containing a 10 bp UMI and a 6 bp sample barcode. To obtain digital exon count matrices, we first used functions `bbduk` to filter out reads that have low sequence complexity (estimated entropy*<*0.5) and `repair` to pair the remaining reads from BBTools, BBMap v. 38.02 (Bushnell 2014). Then we applied zUMIs with default parameters (Parekh et al. 2018, version 2.9.7c, STAR v.2.6.2). Human samples were mapped to hg38 with annotations from Gencode v.32. Cynomolgus *Macaca fascicularis* samples were mapped to macFas6 (Jayakumar et al. 2021) and for gene annotation, we transferred human Gencode v.32 gene models to macFas6 using Liftoff v1.6.3 (Shumate and Salzberg 2021). All samples from time points 0 and 1 were rather homogeneous and showed iPSC characteristics, while all later samples were neural progenitor cells (NPCs). Hence for all analyses, we refer to NPCs as the time points 5, 7, 9. The counts were filtered for UMI counts in at least 28.57% of NPCs (6/21 samples), resulting in a set of 14,608 genes.

### ATAC-seq data generation and processing

#### Data generation

iPSCs of 2 clones from 2 human individuals and 2 clones of 2 cynomolgus macaque individuals were differentiated using the protocol as described above. The NPC lines were cultured in NPC proliferation medium and passaged 2 - 4 times until they were dissociated and subjected to ATAC-seq together with the respective iPSC clones.

ATAC-seq libraries were generated using the Omni-ATAC protocol (Corces et al. 2017) with minor modifications. In brief, cells were washed with PBS and dissociated using Accumax (Sigma Aldrich) for iPSCs or TrypleSelect (Thermo Fisher) for NPCs at 37°C for 5 - 10 min. After cells were counted, 100,000 cells were pelleted at 500 rcf for 5 min, washed with 1 ml PBS and pelleted at 500 rcf for 5 min at 4 °C. The supernatant was removed completely and cells were resuspended in 100 μl chilled nuclei lysis buffer (10 mM Tris-HCl pH7.4, 10 mM NaCl, 3 mM MgCl2 in water, supplemented with 0.1% Tween-20, 0.1% NP40, 0.01% Digitonin and 1% BSA) by pipetting up and down three times, followed by incubation on ice for 3 min. After lysis, 1 ml of lysis wash buffer (10 mM Tris-HCl pH7.4, 10 mM NaCl, 3 mM MgCl2 in water, supplemented with 0.1% Tween-20 and 1% BSA) was added, and tubes were inverted three times. After counting, 50,000 nuclei were pelleted at 500 rcf for 10 min at 4°C, the supernatant was removed and nuclei were resuspended in 50 μl transposition mix (25 μl 2x TD buffer, 2.5 μl TDE1, 16.5 μl PBS, 0.5 μl 1% digitonin, 0.5 μl 10% Tween-20 and 5 μl ddH2O) by pipetting six times. Transposition reactions were incubated at 37 °C for 1 h at 1000 rpm shaking, followed by a clean-up using the DNA Clean & Concentrator-5 kit (Zymo). For library generation, 20 μl of the transposed sample was mixed with 2.5 μl 25μl p5 custom primer, 2.5 μl 25μl p7 custom primer (Buenrostro et al. 2013) and 25 μl NEBNext Ultra II Q5 2x Master Mix (NEB) and a PCR with 10 cycles was conducted as stated in the Omni-ATAC protocol. Libraries were purified using the DNA Clean & Concentrator-5 kit, run on a 2% E-Gel (Thermo Fisher) and gel excision of DNA between 150 bp and 1,500 bp was performed using the Monarch DNA Gel Excision Kit (NEB). Concentrations of the purified libraries were measured using PicoGreen (Thermo Fisher) and quality was assessed using a Bioanalyzer High-Sensitivity DNA Analysis Kit (Agilent). Libraries were pooled and sequenced on NovaSeq 6000 instrument with the following setup: R1: 151, i7: 8, R2: 151 cycles.

#### Data processing

Sequenced human and cynomolgus macaque reads were mapped to hg38 and macFas6 genomes, respectively. For mapping, we used bwa-mem2 (Vasimuddin et al. 2019, version 2.0pre2), using the following command: `bwa-mem2 mem -M -t 20 -I 250,150`. Furthermore, `samtools fixmate -m - -` and `samtools sort` commands were applied (H Li and Durbin 2009, version 1.11). Peak calling was performed using Genrich (https://github.com/jsh58/Genrich) on the 2 biological replicates per species per cell type. We applied the following parameter settings: `-j -y -r -q 0.05 -a 200 -e MT,Y -E $blacklist -s 20`, where as a `$blacklist` the ENCODE blacklist with hg38 coordinates (Amemiya et al. 2019) was supplied for human (910 regions), and a reciprocal lift-over version of it to macFas6 (558 regions) was supplied for the peak-calling in macFas6 genome space.

#### Lift-over file generation and usage

Lift-over files hg19toHg38 and hg38ToHg19 were downloaded from USCS (https://hgdownload.soe.ucsc.edu/gbdb/hg19/liftOver/hg19ToHg38.over.chain.gz, https://hgdownload.soe.ucsc.edu/goldenPath/hg38/liftOver/hg38ToHg19.over.chain.gz). Lift-over files hg38toMacFas6 and macFas6toHg38 were generated from blastz alignments (Schwartz et al. 2003; Kent et al. 2003) of the canonical chromosomes from both genomes, as reported here(https://genomewiki.ucsc.edu/index.php?title=DoBlastzChainNet.pl). Reciprocal lift-over (RLO) was used to lift CRE coordinates from hg19 over to hg38 and from hg38 over to macFas6. In both cases, the coordinates from X were lifted to Y, then the matches in Y that carried the same CRE identifier and were *<* 40bp distant from each other were merged and lifted back to X. For further analyses, we kept the CREs of which the reciprocal lift-over coordinates in X overlapped the original sequence coordinates in X. We identified RLO matches for 99.7% of the CREs in hg38 and 87.1% in macFas6. We further removed CREs of which the RLO match width was beyond the following boundaries: [1.2 x hg19; 0.8 x hg19]. We also removed 36 of the remaining CREs that contained Ns in the sequence of either species genome. This resulted in an orthologous set containing 401,389 CREs.

#### Cross-species accessibility and gene expression analysis

ATAC-seq reads from cynomolgus macaque NPCs mapping to macFas6 and from human NPCs mapping to hg38 genomes were counted within the lift-overed PD-CRE coordinates. Only CREs that overlapped with an ATAC-seq peak by 10% relative to the width of both the DHS and the ATAC-seq peak in at least one species were kept for differential accessibility (DA) analysis (n=61,379). Differential gene expression (DGE) and accessibility (DA) analyses were both performed separately using DESeq2 (Love et al. 2014, version 1.38.3), using species as the predictor. A significance level of Benjamini-Hochberg adjusted *p*−value of 0.1 was used to detect DA or DGE.

For further downstream analysis where we used the state of the ATAC-seq peak (open / closed) as the indicator for peak conservation, we furthermore required that conserved peaks need to overlap by 10% of their width between the lifted human peaks to macaque genome and the macaque peaks. In addition, we excluded cases where, in either species, multiple ATAC-seq peaks overlapped the same DHS or vice versa to avoid multi-to-1 and 1-to-multi peak overlaps, leaving us with a set of 1-to-1, 0-to-1, 1-to-0 and 0-to-0 overlaps between DHS-CREs and ATAC-seq peaks in either species.

#### Evolutionary sequence analysis of CREs

To be able to intepret evolution rates as a result of the genetic element’s CRE activity, we excluded all CREs that overlapped CDSs (Gencode v.19) in all sequence evolution analyses (6.6% of the gene-assigned CREs).

##### INSIGHT

We ran the web tool INSIGHT (Gronau et al. 2013) on the CRE or peak coordinates of each PD class in hg19 using the default settings. To re-calculate the evolutionary rates on various CRE subsets more efficiently, we downloaded the INSIGHT script `runINSIGHT-EM.sh` that applies expectation-maximization (EM) algorithm on the provided INSIGHT files (.ins) and the complementary flanking sequence INSIGHT files (`.flankPoly.forBetas.ins`). The scripts for subsetting the INSIGHT output files and re-calculating the evolutionary rates can be found on github.

##### phastCons and phyloP

Pre-calculated 46-way hg19 phastCons and phyloP scores for the 10 primate subset were downloaded from http://hgdownload.cse.ucsc.edu/goldenpath/hg19/phastCons46way/ and http://hgdownload.cse.ucsc.edu/goldenpath/hg19/phyloP46way/ (versions from 2009-11-11) in a big-wig file format. For each CRE, the average conservation score was calculated for each conservation metric.

#### Quantification of transcription factor binding

Two sets of TF Position Weight Matrices (PWMs) of the 1) expressed TFs in 9 tissues from Epigenomics Roadmap project (Bernstein et al. 2010) (643 motifs from 561 TFs) and 2) expressed TFs in our human and cynomolgus macaque NPCs (521 motifs from 446 TFs) were generated by downloading and subsetting JASPAR 2020 collection, core vertebrate set (Fornes et al. 2020) using R packages `JASPAR2020` (version 0.99.10) and `TFBSTools` (Tan and Lenhard 2016, version 1.36.0). These PWMs were provided to Cluster-buster (Frith et al. 2003, downloaded on 2020-05-07). Cluster-Buster was ran on each set with the following settings: `-c0 -m0 -r10000 -b500 -f5`. The orthologous human and cynomolgus macaque CRE input sequences were extended by 500 bp in each direction, allowing cluster-buster to have a better approximation of the background base composition (parameter `-b500`). In each species for each TFBS cluster of a CRE, we ranked TF motifs based on their strongest binding site. For all subsequent analyses, for each CRE we only considered TF binding motifs that were among the 10% strongest in at least one cluster in at least one species.

#### TFBS diversity and divergence between human and macaque orthologous CREs

For each CRE in each species, we measured TFBS diversity by Shannon entropy (*H*) (Shannon 1948) where we considered a CRE as a collection of *i* = 1, 2, .., *n* motifs of varying frequency (*p*):

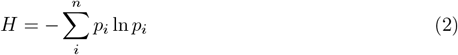

For each CRE, motif scores were estimated for each enriched motif (see Methods section Quantification of transcription factor binding) along the sequence by Cluster-Buster (Frith et al. 2003) and they were used as a proxy for TF binding potential. We summed up the motif scores for each motif to obtain the cumulative motif score. We then used it to calculate *p* in relation to the total cumulative score of all motifs combined. As entropy is an index of diversity instead of diversity itself, *H* was converted to what is known as true diversity or Hill number of order 1 (Hill 1973; Jost 2006) simply by

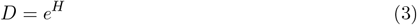

which measures the effective cumulative motif score.

In order to measure how TFBS repertoires diverge between the two species, we calculated the average Canberra distance 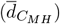 for each CRE across the *i* = 1, 2, .., *n* motif cumulative scores (*S*) as follows:

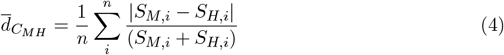

where *M* indicates the orthologous CRE in macaque and *H* in human. Further, we used

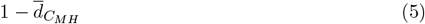

as a proxy for TFBS repertoire conservation.

#### CRE PD ranking per motif to detect over-represented motifs

##### Per tissue

We first identified the expressed TFs and their respective motifs and considered only their binding to the CREs that are open in that tissue. For each PD category and motif, the relative binding frequency was obtained as the fraction of CREs that have binding sites for that motif, e.g.

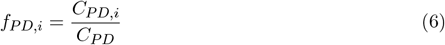

where *PD* indicates a PD category, *i* indicates a motif, *C*_*P D,i*_ is the count of CREs with motif *i* binding site(s) present in the particular *PD* category, *C*_*P D*_ is the total CRE count in that *PD* category. Having obtained these relative frequencies per PD, we then ranked PD categories for each motif. Fold changes of the binding fraction of rank-1 PD relative to the average fraction were calculated for each motif *i* as:

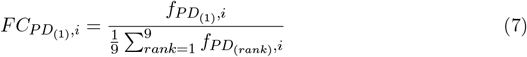

##### Across tissues

To summarise motif-PD enrichment across tissues, we focused on motifs that had the highest binding fractions (rank-1) to either PD9 or PD1. To obtain the PD9-enriched motifs, we identified TF motifs for which PD9 CREs had rank-1 in all tissues. As the PD1-tissue-specific motifs we considered the ones that have PD1 with rank-1 only in that particular tissue, but not in the other tissues. Gene-set enrichment analysis contrasting the respective TF groups with the rest of the expressed TFs was conducted using the Bioconductor package topGO (Alexa and Rahnenfuhrer n.d., version 2.50.0), setting the following parameters: `ontology=“BP”, nodeSize = 10, algorithm = “elim”, statistic = “fisher”` .

#### Stripe factor enrichment analysis

Stripe factor annotation table was obtained from Zhao et al. 2022. We selected the stripe factors detected in human (“Human Stripe Factors”) and to subset the universal stripe factors, we used a cutoff of 0.9 for the proportion of total samples in which this TF was detected to be a stripe factor.

#### TFBS position overlap between human and macaque orthologous CREs

Orthologous human and macaque CRE sequences were pairwise aligned using mafft (Katoh and Standley 2013, version 7) using the following parameters `--adjustdirection --maxiterate 1000 --auto`. We quantified alignment length (median 1273, 90% CI [1133, 1790]), fraction of mismatches in bp (median: 0.058, 90% CI [0.0357, 0.1432]), the fraction of indels in bp (median: 0.018, 90% CI [0.0031, 0.0916]) and the number of indels (median 6, 90% CI [2, 13]). We subsequently trimmed gaps in the remaining CRE alignments.

Using the alignment of a CRE, the positions of TFBS that had a motif binding score of *>*=3 in either species were projected onto the common alignment space. Binding site agreement per motif *i* was calculated as the intersection of binding positions in bp between species over the union, also known as Jaccard similarity coefficient, and summarized by taking the mean across all *i* = 1, 2, .., *n* motifs that bind to the particular CRE:

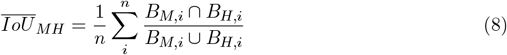

where *B* is a set of positions in the alignment that overlap with a binding site of motif *i* in the respective species macaque *M* or human *H*.

#### Quantification and Statistical Analysis

Data visualizations and statistical analysis was performed using R (version 4.2.3) (R Core Team 2023), session info can be accessed on GitHub. Details of the statistical tests performed in this study can be found in the main text as well as the method details section. Schematics were made using bioRender.

## Data and Code Access

RNA-seq and ATAC-seq data are available under ArrayExpress accessions E-MTAB-13494 and E-MTAB-13373. A compendium containing processing scripts, important tables and detailed instructions to reproduce the analysis for this manuscript is available from the following GitHub repository: https://github.com/Hellmann-Lab/The-effects-of-pleiotropy-on-regulatory-evolution Data files and tables are deposited on zenodo (DOI: 10.5281/zenodo.10471368).

## Acknowledgements

We thank Lucas Esteban Wange, Johannes Bagnoli, Aleks Janjic for helping with bulk RNA-seq preparation. We also thank Paulina Spurk for contributing to the initial ATAC-seq analysis, Diego Emiliano Ruiz Navarro for contributing to the initial RNA-seq analysis, Swati Parekh for assistance with perl scripts and Eva Briem for helping with a schematic. We also thank Boyan Bonev for generating the liftOver files from hg38 to macFas6 and Andrea Betancourt for helpful discussions.

## Author Contributions

I.H. proposed the project and conceived the approaches of this study. W.E. provided the resources for data generation and helpful discussions. P.O. processed the human tissue accessibility data. B.V. provided expertise during initial steps. V.Y.K.L. contributed to TFBS evolutionary analyses. J.G. and M.H generated the primate cell lines and the expression data. S.K. and J.G. generated the primate accessibility data. I.H. supervised the work and provided guidance in data analysis. Z.K. collected, integrated and analysed all data. Z.K. and I.H. wrote the manuscript. All authors read, corrected and approved the final manuscript.

## Supplemental Information

**Supplemental Figure S1.**
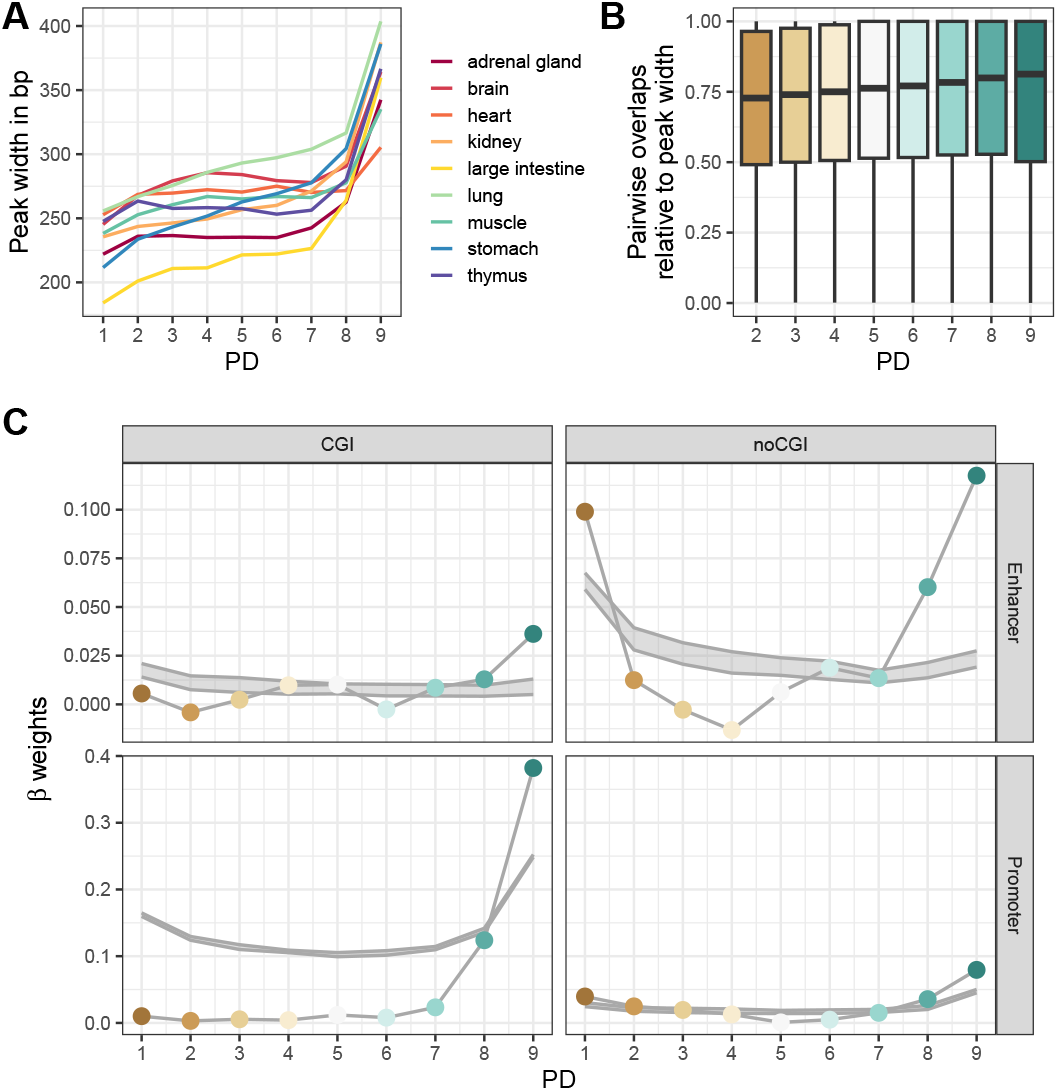
Peak widths per tissue and overlaps across tissues. **A**. Average peak width per tissue across specificity groups prior to cross-tissue merging. **B**. Pairwise overlap fraction between overlapping peaks that were later merged into the same CRE. **C**. Observed coefficients for the CREs from different PDs (thin line with colored points) are different from the control estimates (90% CI, in gray) where PD labels were shuffled 30 times across the CREs.

**Supplemental Figure S2.**
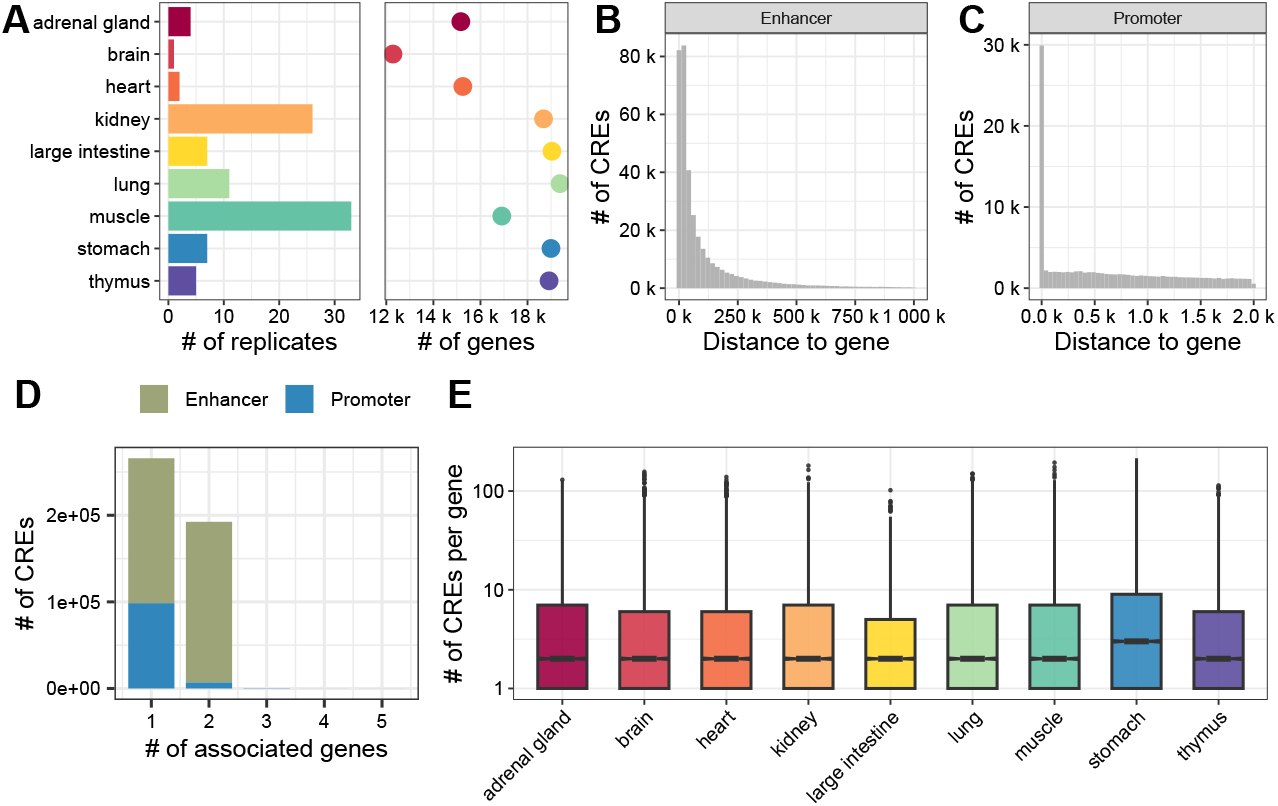
CRE to gene association across tissues. **A**. Expression data overview: Number of replicates and number of expressed genes for each tissue. **B-C**. Distance distribution of CREs annotated as enhancers (**B**) and promoters (**C**) to their closest gene. **D**. Number of associated genes per CRE. Enhancers were associated with up to 2 genes, promoters: up to 5. **E**. Associated number of CREs per gene is comparable across tissues.

**Supplemental Figure S3.**
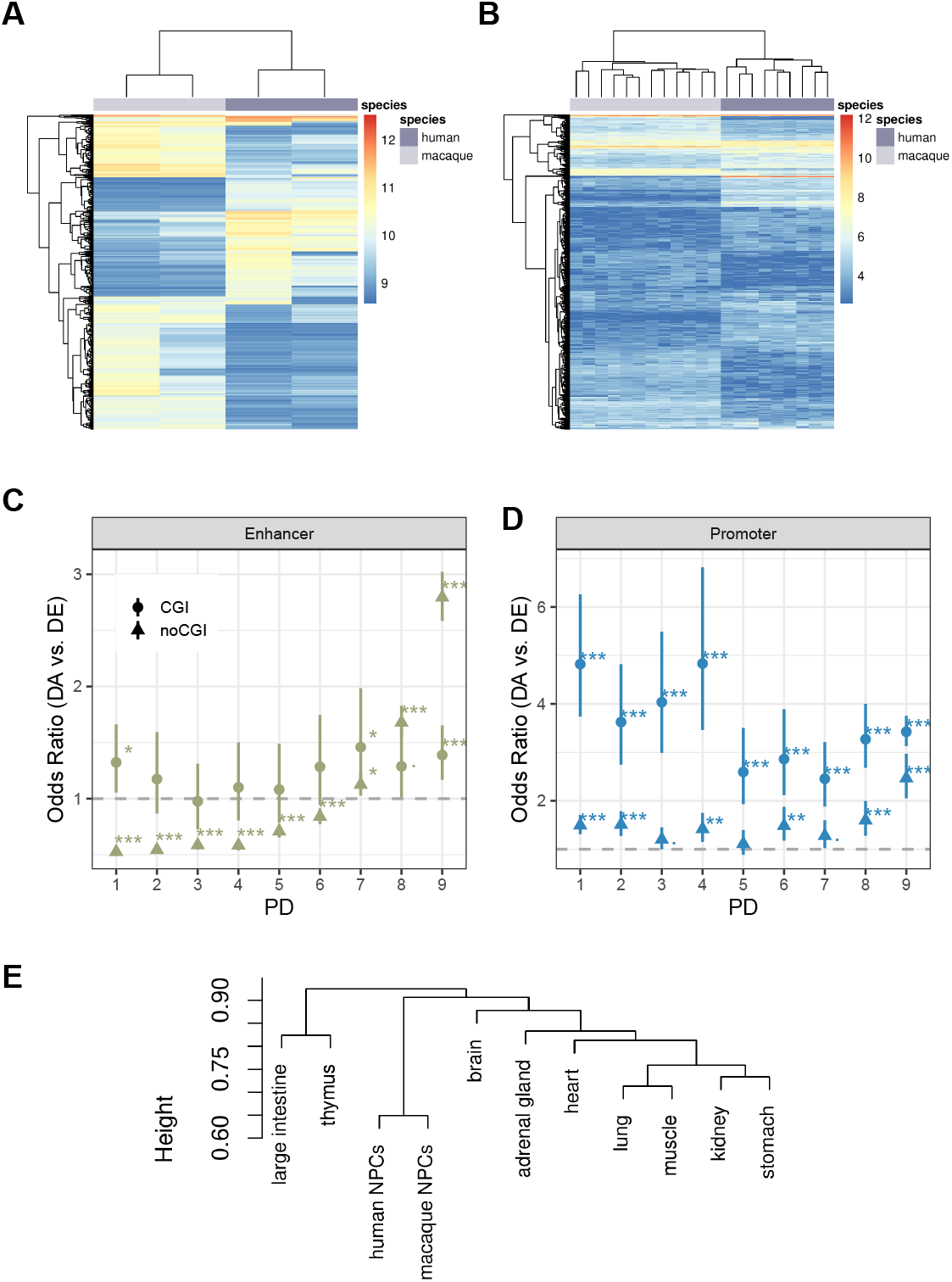
Cross-species NPC expression and accessibility. **A**. Heatmap of the top 1000 most variable CREs across the ATAC-seq data in terms of their openness (Euclidean distance, method: complete). **B**. Heatmap of the top 1000 most variable genes across the RNA-seq data (Euclidean distance, method: complete). **C**. Odds ratios of differential accessiblity of enhancers vs. the associated gene differential expression results between humans and cynomolgus macaques, split between CGI and non-CGI CREs. **D**. Odds ratios of differential accessiblity of promoters vs. the associated gene differential expression results between humans and cynomolgus macaques, split between CGI and non-CGI CREs. **C**,**D**. Error bars represent the 95% confidence intervals of the odd ratio, the stars indicate the significance level after Benjamini-Hochberg correction ( . *<* 0.1, ^*^ *<* 0.05, ^**^ *<* 0.01, ^***^ *<* 0.001 ). **E**. Hierarchical clustering (method = “binary”, distance = “complete”) of human and macaque NPC ATAC-seq samples together with human tissue CREs based on the variable CRE binary openness across tissues (PD1-8).

**Supplemental Figure S4.**
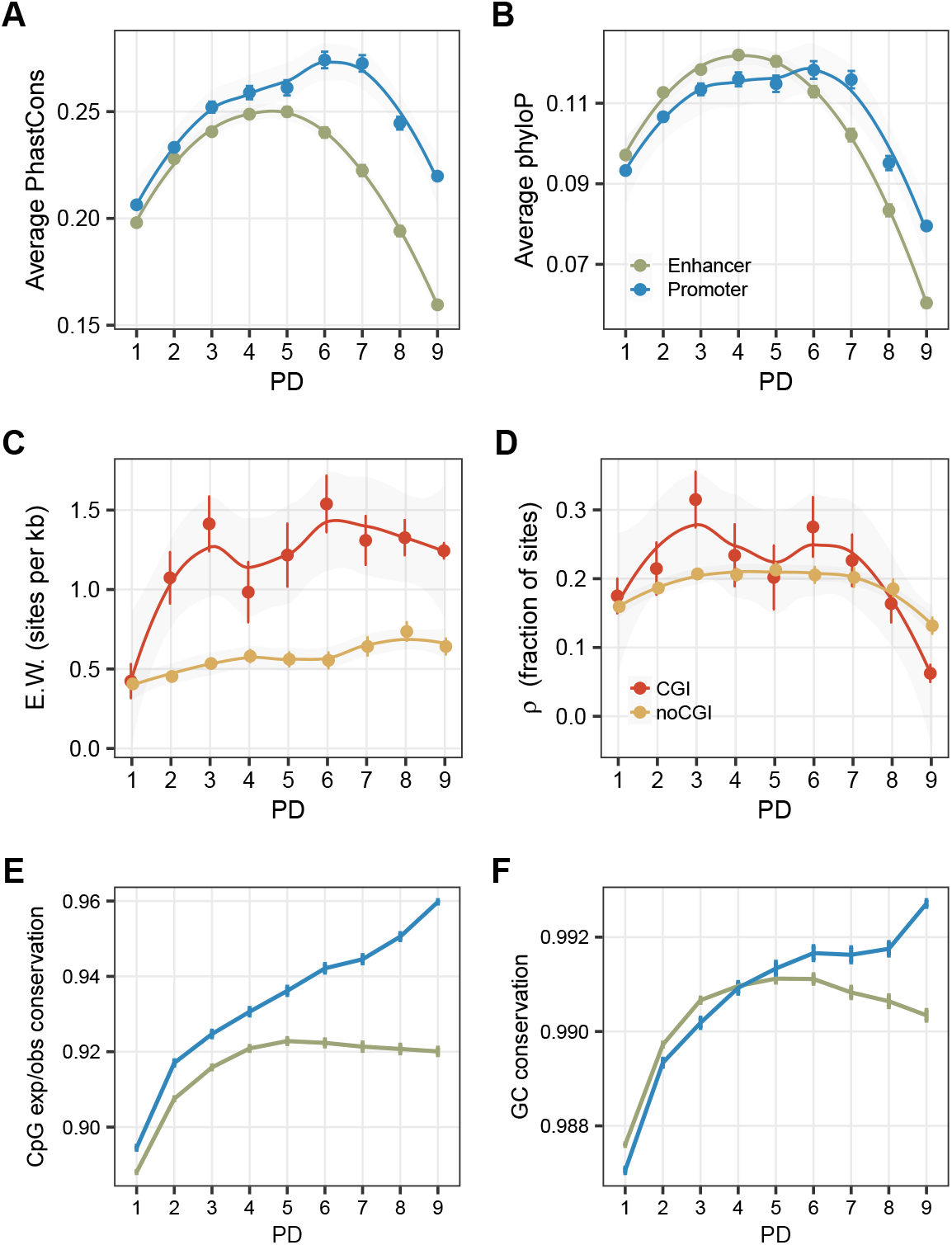
Evolutionary sequence analysis of CREs across tissue specificity groups. **A**. PhastCons conservation scores based on a 10-species primate tree. **B**. PhyloP conservation scores based on a 10-species primate tree. **C**. Weak negative selection patterns between CGI and non-CGI CREs suggest an increasing weak selection with higher PD in both groups. **D**. (Strong) negative selection patterns between CGI and non-CGI CREs show similar trend across PDs. **E**. Absolute pairwise distance in CpG expected/observed ratio between human and cynomolgus macaque orthologous CREs. **F**. Absolute pairwise distance in GC content between human and cynomolgus macaque orthologous CREs. **A**,**B**,**C**,**D**,**E**,**F**. Depicted are mean estimates per PD category. Error bars indicate SEM.

**Supplemental Figure S5.**
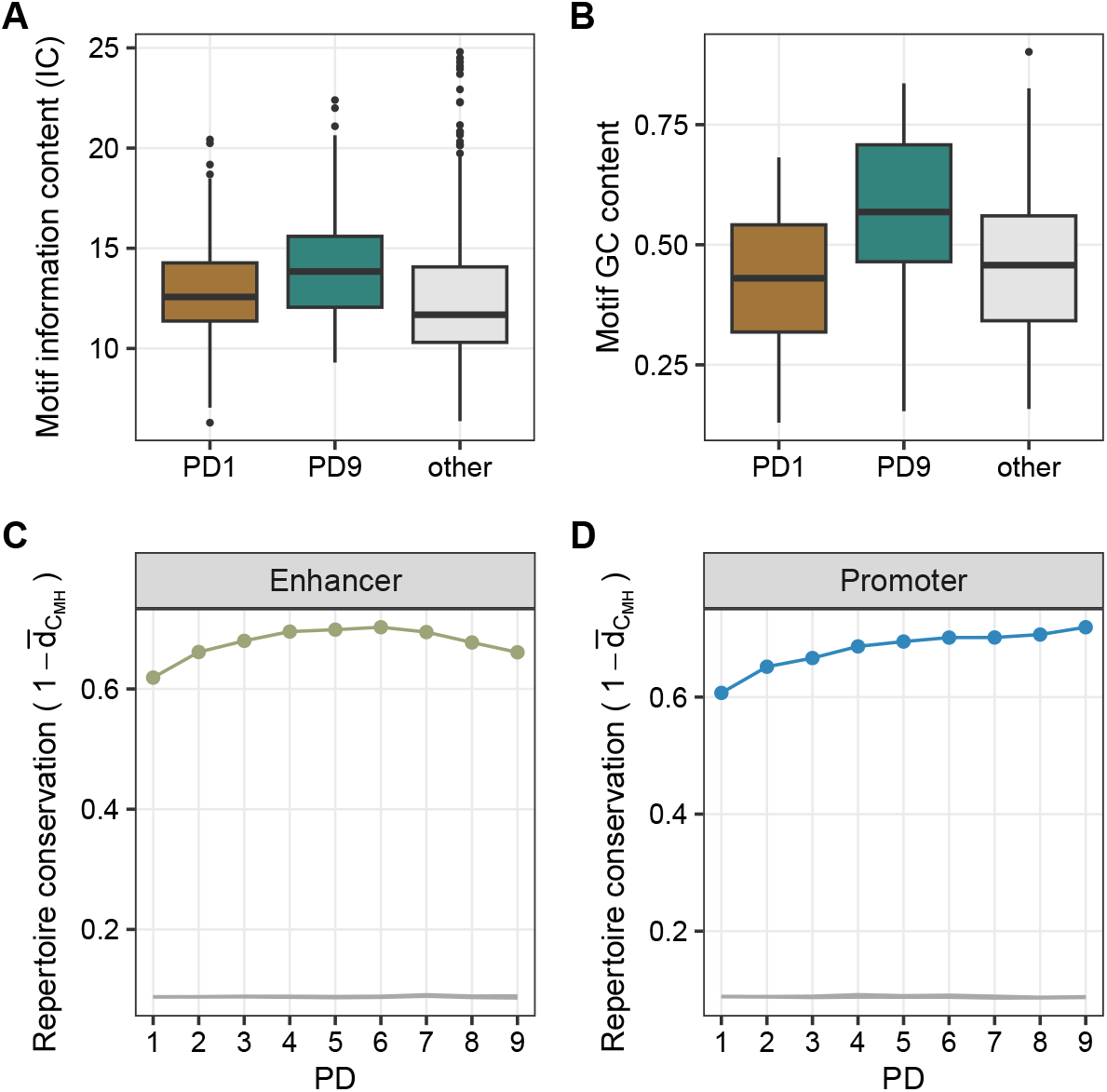
TFBS repertoire diversity and conservation. **A**. Motif information content of the PD1-enriched, PD9-enriched and other motifs, classified as in 2D. **B**. Motif GC content of the PD1-enriched, PD9-enriched and other motifs, classified as in 2D. **C**,**D**. 95% confidence intervals of the average TFBS repertoire conservation when shuffling macaque CRE identifiers within the respective PD class 10 times (grey line). Random CRE similarity is below 10% and does not increase with PD. In comparison, the real observed enhancer and promoter repertoire conservation is depicted in green (**C**.) and blue (**D**.), respectively.

